# Synthetic Poly(2-Oxazoline) Chimera “Lipo-Polymers” as Replacements for Key Lipids in Lipid Nanoparticles

**DOI:** 10.64898/2026.10.23.761445

**Authors:** Jacob D. Ramsey, Matthew J. Haney, Yuling Zhao, Hallie Hutsell, Alexander V. Kabanov

## Abstract

Lipid nanoparticles (LNPs) rely on specialized lipid components, many of which are proprietary and difficult to access or modify. Here, we introduce a polymer-based alternative to key LNP constituents through the development of chimera poly(2-oxazoline) (POx) synthetic lipid polymers (SLPs) and synthetic lipid oligomers (SLOs). These chimeric lipid-mimetic materials are designed for seamless incorporation into existing LNP architectures, enabling direct replacement of PEG-lipid conjugates and ionizable lipids, respectively. Chimeric SLPs comprise hydrophilic shell-forming POx blocks linked to hydrophobic lipid-inserting domains, whereas SLOs consist of hydrophobic POx oligomers terminated with ionizable amine-containing groups. Using mRNA-loaded LNPs, we demonstrate that first-generation POx-Chimera SLPs and SLOs functionally replace industry-standard lipid components while maintaining robust transfection activity *in vitro* and *in vivo*. Notably, SLO-containing formulations achieved expression levels comparable to, and in some cases exceeding, those of conventional lipid formulations. Together, these results establish POx-Chimera SLPs and SLOs as a versatile platform for polymer-centric LNP engineering, creating new opportunities to tune nanoparticle composition, biodistribution, and transgene expression while reducing reliance on proprietary lipids.

## TEXT

Following their immense success as a vaccine platform during the COVID-19 pandemic, lipid nanoparticles (LNPs) have become the industry-standard drug delivery system for nucleic acid cargoes. Encapsulation of nucleic acids in LNPs relies on four core lipid components: ionizable lipids, cholesterol, helper-lipids, and a poly(ethylene-glycol) (PEG) lipids^1–4^. Significant efforts over the past several years have focused on modulating LNP biodistribution and organ-specific accumulation. Changes in lipid chemistry alter LNP physicochemical properties and structural morphology, enabling preferential targeting of specific organs in preclinical settings, including lung, spleen, and other tissues^5–9^. In recent work, we have modulated LNP distribution and expression profiles via conjugation of poly(2-oxazoline) (POx) polymers to lipids, replacing the conventional PEG-lipid component^10^. We found that shorter POx lipids increased peak muscle expression, enhanced expression durability, and altered distribution to select lymphoid organs. POx–lipids also improved transfection efficiency in dendritic cells, macrophages, and tumor cells while reducing transfection in hepatic cells *in vitro* and decreasing liver tropism *in vivo*^10^.

Beyond their role in modulating *in vivo* expression profiles, POx polymers serve as a compelling alternative to PEG-lipids, independent of their effects on biodistribution and expression^11–16^. The widespread presence of anti-PEG antibodies in the general population necessitates identification of viable alternatives. Continued use of PEGylated nanoparticles risks increased anaphylactoid responses and reduced efficacy due to accelerated blood clearance of PEG-containing products^17–19^. Accordingly, alternative polymers lacking prevalent pre-existing antibodies are needed to mitigate these effects^20,21^.

In this work, we introduce a new class of POx-based synthetic lipid polymers (SLPs) as an alternative to conventional POx and PEG–lipid conjugates. These chimera lipo-polymers establish a fully polymer-based platform for LNP formulation, eliminating the need for proprietary lipids. We further introduce POx-based ionizable synthetic lipid oligomers (SLOs) as a replacement for conventional ionizable lipids. Compared with traditional lipid synthesis, POx-Chimera SLP and SLO manufacturing is simple, modular, scalable, and potentially more cost-effective. Furthermore, the approach avoids post-synthetic conjugation and eliminates the need for linker chemistries.

We have synthesized the first generation of POx-Chimera SLPs using a di-block architecture comprising a hydrophilic poly(2-methyl-2-oxazoline) (PMeOx) block and a hydrophobic, lipid-mimetic poly(2-nonyl-2-oxazoline) (PNOx) oligo-block. Together, these two blocks mimic the structure of a POx–lipid conjugate and can be readily incorporated into established LNP formulations (**Figure 1A**). The PMeOx block was designed to have a molecular weight of approximately 2,000 g/mol, comparable to that of commonly used POx and PEG lipids. This polymer was designated **PMeOx-(NOx)ₙ**. The polymer was terminated with a piperidine group, rendering it functionally an ionizable lipo-polymer.

**Figure 1:**
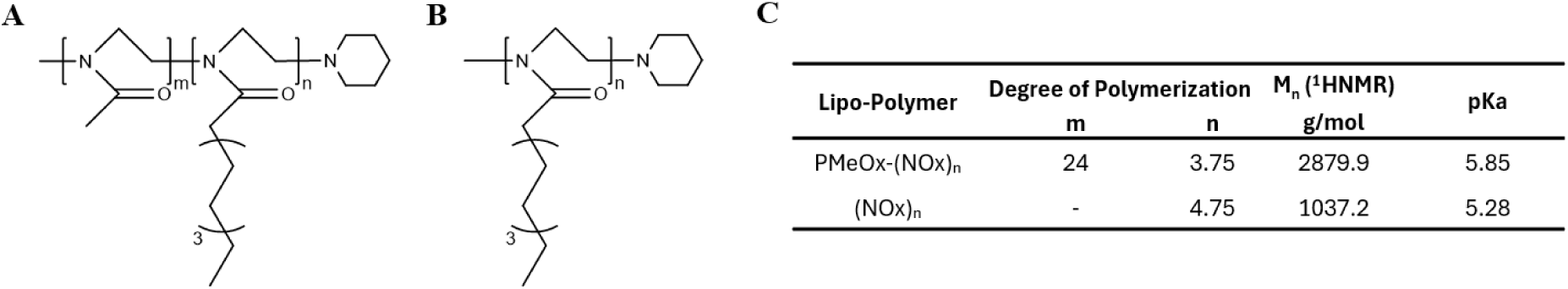
Structure and characterization of Generation 1 POx-Chimera lipo-polymers. Chemical structures of (A) PMeOx-(NOx)_n_ and (B) (NOx)_n_ with (C) ^1^HNMR determined block lengths, molecular masses (M_n_), and pKa determined by titration.

We further hypothesized that removal of the PMeOx block would yield a SLO capable of directly replacing an ionizable lipid. This SLO consists solely of the lipid-mimetic **(NOx)ₙ** oligomer and a terminal ionizable piperidine group (**Figure 1B**). **Figure 1C** summarizes the block lengths and molecular masses (M_n_) determined by ^1^HNMR for the synthesized polymer and oligomer. ^1^HNMR spectra for all polymers and oligomers, as well as gel permeation chromatography (GPC) elution profiles and dispersity data for PMeOx-(NOx)ₙ are in **Supplemental Figures 1-3**. The SLO (NOx)ₙ was too hydrophobic for reliable GPC analysis, exhibiting strong interactions with the column stationary phase that precluded accurate determination of its molecular weight by this method.

To assess the ionization behavior of the SLP and SLO, potentiometric titrations were performed to determine their pKa values. The pKa of PMeOx-(NOx)ₙ and (NOx)ₙ to be 5.85 and 5.28, respectively (**Figure 1C**). These values are comparable to those reported for the ionizable lipids, including clinically validated SM-102 and ALC-0315 lipids (6.68 and 6.09, respectively; Broadpharm® datasheets), supporting the use of (NOx)ₙ as a potential substitute for conventional ionizable lipids in LNP formulations when formulated at pH 4.0. The titration curves and first derivatives of the titration curves used to determine equivalence points and pKa are show in **Supplemental Figure 4.**

We next optimized PMeOx-(NOx)ₙ as a direct replacement for DMG-PEG in a standard LNP formulation containing DMG-PEG and ALC-0315 (Pfizer’s ionizable lipid). To identify an appropriate PMeOx-(NOx)ₙ content (mol %), we prepared small batches of firefly luciferase (fLuc) mRNA mRNA-loaded LNPs using a modified high-throughput shaker method^22^.

Formulations containing 5 μg of mRNA were characterized for particle stability by dynamic light scattering (DLS) over four days and for mRNA encapsulation efficiency (**Figure 2A**). DLS measurements are summarized in **Supplemental Table 1**, while encapsulation efficiencies determined by the RiboGreen assay are shown in **Figure 2B**.

**Figure 2.**
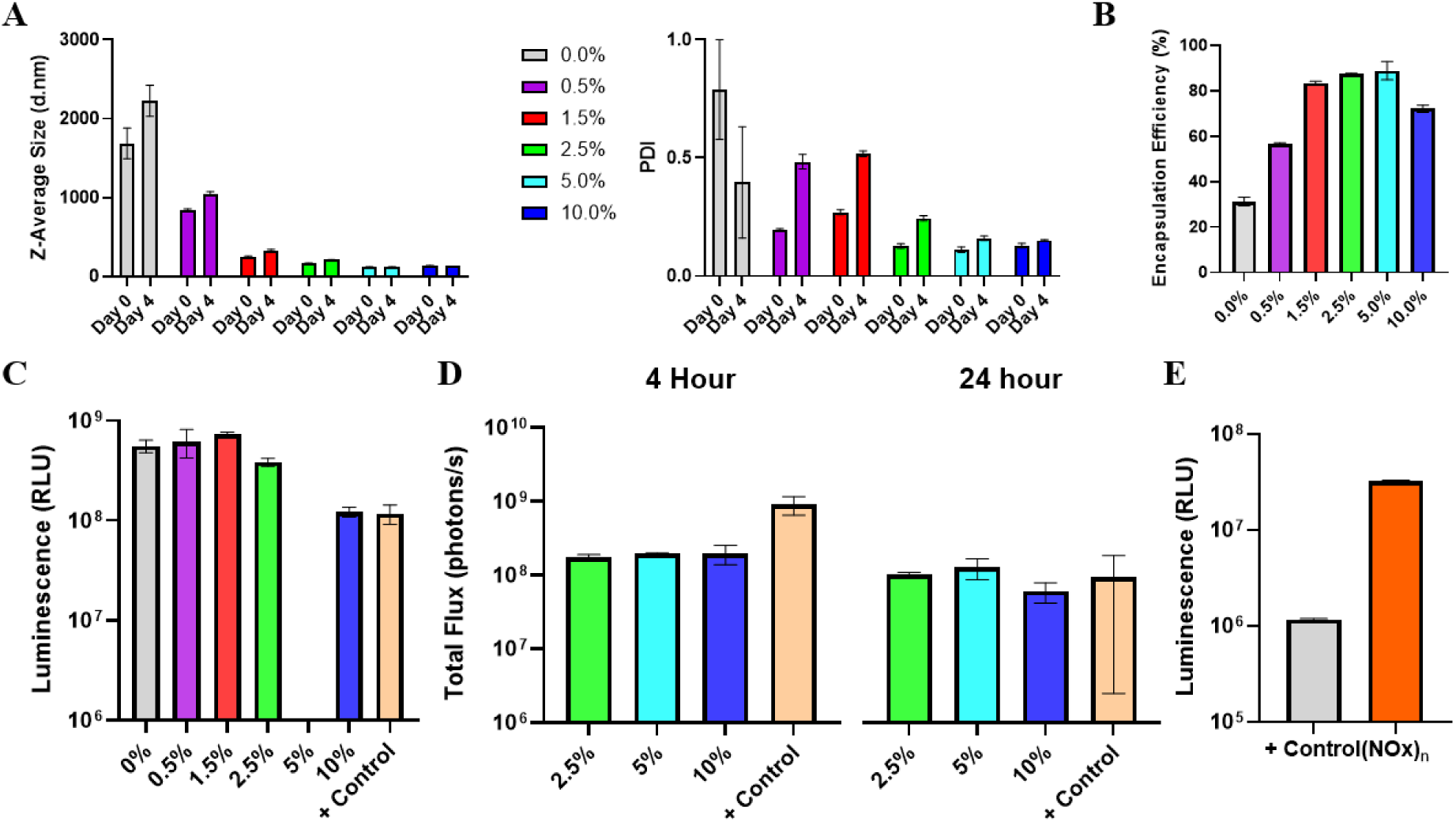
Stability and functional activity of POx-based synthetic lipid-mimetic polymer and oligomer formulations. (A) DLS-measured particle size and PDI stability during 4 days of storage at 4°C. (B) RiboGreen-determined mRNA encapsulation efficiency. (C) *In vitro* transfection of HEK293 cells. (D) *In vivo* transfection following IM administration (N=2/group). (E) *In vitro* transfection of HEK293 cells with LNPs formulated using (NOx)ₙ as an ionizable lipid substitute, in which ALC-0315 was replaced by (NOx)ₙ at 50 mol%. For panels A–D, the mol% of PMeOx-(NOx)ₙ in the LNP formulation is indicated. For all panels, the positive control was the standard LNP formulation containing 50 mol% ALC-0315, 1.5 mol% DMG-PEG, 10 mol% DSPC, and 38.5 mol% cholesterol.

The base formulation consisted of 50% ionizable lipid, 1.5% polymer-lipid, 10% DSPC, and 38.5% cholesterol. To increase the content of PMeOx-(NOx)ₙ from 0 to 10% mol%, equivalent amounts of DSPC and cholesterol were removed from the formulation. As shown in **Figure 2C**, PMeOx-(NOx)ₙ-containing formulations achieved *in vitro* transfection efficiencies in HEK293 cells comparable to those of the ALC-0315/DMG-PEG control (a technical error in the 5% group resulted in null data). Because only formulations containing ≥2.5 mol % PMeOx-(NOx)ₙ remained stable over four days and exhibited high encapsulation efficiency, these formulations were further evaluated following intramuscular (IM) administration in BALB/c mice (**Figure 2D**). PMeOx-(NOx)ₙ containing LNPs produced approximately 0.5–1 log lower luminescence than the control formulation at 4 hours post-injection but achieved comparable expression by 24 h. Finally, in a separate experiment, we confirmed that the SLO (NOx)ₙ could also function as a direct replacement (50 mol% ) for ALC-0315 in LNP formulations along with DMG-PEG (**Figure 2E**).

We selected the 5 mol % PMeOx-(NOx)ₙ formulation for further studies. Larger batches containing 50 μg of fLuc mRNA were then prepared by impinged jet mixing using four formulations: (i) ALC-0315 with DMG-PEG, (ii) ALC-0315 with PMeOx-(NOx)ₙ, (iii) (NOx)ₙ with DMG-PEG, and (iv) (NOx)ₙ with PMeOx-(NOx)ₙ. In formulations containing both replacements, (NOx)ₙ substituted for ALC-0315 on a 1:1 molar basis, while PMeOx-(NOx)ₙ replaced DMG-PEG at 5 mol%, with DSPC and cholesterol each reduced by 1.75 mol% to maintain the overall formulation composition (See **Supplemental Table 2** for compositions). Particle size and PDI were characterized by DLS, and mRNA encapsulation efficiency was determined using the RiboGreen assay (**Table 1**). All formulations exhibited comparable particle sizes, narrow size distributions, and high encapsulation efficiencies.

**Table 1.** Particle size, PDI, and mRNA encapsulation efficiency of LNP formulations measured by DLS and the RiboGreen assay.

| Lipid Composition |  | Z-Average Size | PDI | Encapsulation Efficiency |
| --- | --- | --- | --- | --- |
| Ionizable Lipid/SLO | Polymer Lipid/SLP | (nm) | - | (%) |
| ALC-0315 | DMG-PEG | 69.5 ± 0.3 | 0.206 ± 0.002 | 85.2 ± 3.94 |
| ALC-0315 | PMeOx-L | 63.9 ± 0.5 | 0.208 ± 0.010 | 77.5 ± 2.19 |
| PNOx-L | DMG-PEG | 61.0 ± 0.3 | 0.089 ± 0.011 | 93.3 ± 3.23 |
| PNOx-L | PMeOx-L | 53.7 ± 0.1 | 0.078 ± 0.001 | 92.2 ± 2.84 |

Each LNP formulation prepared by impinged jet mixing was administered IM to BALB/c mice (N=5 per group), a clinically relevant route for vaccine delivery. Luciferase expression was monitored for 72 hrs following administration (**Figure 3A** and **B**). The control formulation containing ALC-0315 and DMG-PEG produced the highest initial level of muscle expression, which declined steadily over 4-72 hours. Replacement of DMG-PEG with PMeOx-(NOx)ₙ resulted in the lowest muscle expression at all time points; however, expression remained within approximately one order of magnitude of the control formulation containing commercial lipids, suggesting substantial opportunity for further optimization. In contrast, formulations containing the ionizable SLO (NOx)ₙ exhibited remarkably stable muscle expression from 4 to 48 hrs before declining at 72 hrs. Analysis of liver signal revealed the formulation containing ALC-0315 consistently produced the highest hepatic expression relative to muscle (**Figure 3G**). Formulations containing (NOx)_n_ showed markedly lower levels of hepatic expression. These results suggest that the identity of the ionizable lipid/lipo-polymer influences hepatic transgene expression and biodistribution, while the (NOx)ₙ SLO can support sustained expression in muscle.

**Figure 3.**
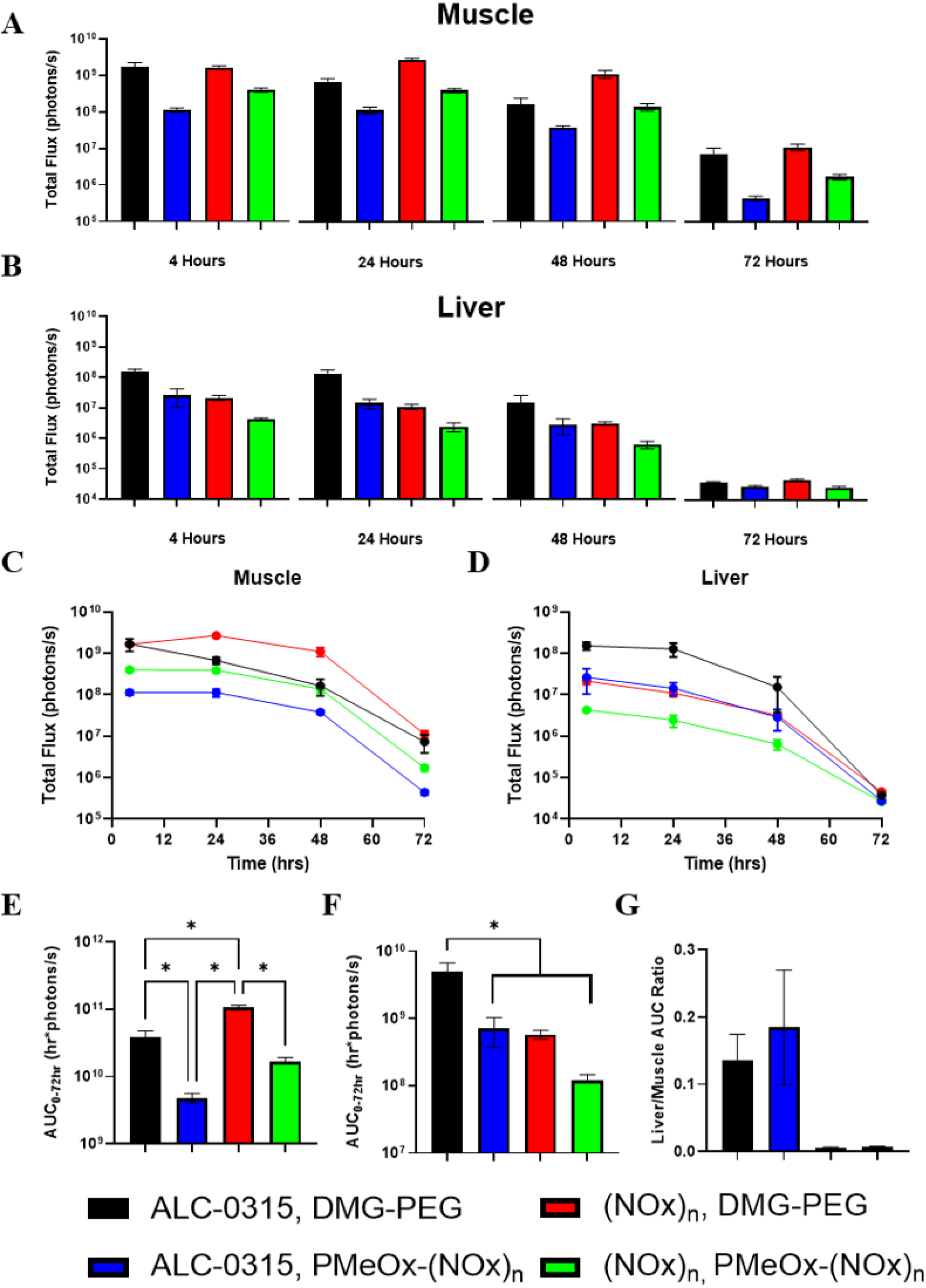
*In vivo* transfection and durability of expression profiles comparing novel lipo-polymer systems to LNPs formulated with industry-standard formulation. Expression of fLuc mRNA measured by IVIS for each formulation at each time point (4, 24, 48, and 72 hours) in **(A)** the muscle and **(B)** the liver. Full-time courses of each formulation in **(C)** the muscle and **(D)** the liver. AUC from 0-72 hrs for **(E)** the muscle and **(F)** the liver. **(G)** Ratio of liver to muscle AUC. N=5 mice/group for all time points.

Complete expression time courses are presented in **Figure 3C,D**, while the corresponding data plotted on a linear y-axis are provided in **Supplemental Figure S5**. The areas under the curve (AUC) for the muscle and liver expression are summarized in **Figures 3E** and **F**. The ionizable SLO (NOx)ₙ increased muscle AUC by approximately one order of magnitude relative to the formulation containing commercial lipids. Although direct replacement of DMG-PEG with PMeOx-(NOx)ₙ resulted in the lowest muscle AUC, incorporation of (NOx)ₙ into this formulation restored expression to levels comparable to those of the industry-standard formulation. Notably, the synthetic PMeOx-(NOx)ₙ/(NOx)ₙ formulation exhibited the lowest liver accumulation, whereas the formulation containing ALC-0315 and DMG-PEG showed the highest. Comparison of liver-to-muscle AUC ratios further revealed that (NOx)ₙ co-formulated with either DMG-PEG or PMeOx-(NOx)ₙ had the lowest values, highlighting the ability of the ionizable SLO (NOx)ₙ reduce off-target liver expression while maintaining robust expression at the IM injection site.

There is a clear need for PEG-alternatives that address emerging challenges in modern medicine. The work of our group and others has helped position POx as a leading alternative to PEG as a stabilizing polymer in LNP formulations^10–16^. In the present work, we describe a fundamentally different approach to POx-based LNP design through the development of POx SLPs as replacement to POx-lipid conjugate and SLOs as replacement for conventional ionizable lipids. POx-Chimera SLPs and SLOs offer several potential advantages, including enhanced structural tunability, simplified synthesis, and reduced manufacturing cost. A unique strength of the POx platform is the ability to synthesize well-defined block copolymers with precise control over composition and architecture. Combined with the broad range of available 2-oxazoline monomers, this versatility provides extensive opportunities for molecular and formulation optimization, enabling the rational tuning of biodistribution, expression kinetics, and other performance characteristics.

The present study demonstrates the capacity of POx SLPs and SLOs to directly and functionally replace industry-standard lipid components in LNP formulations. The SLP PMeOx-(NOx)ₙ performed well *in vitro* as a direct replacement for DMG-PEG. Although *in vivo* transfection levels were approximately one order of magnitude lower than those achieved with the industry-standard formulation, the di-block copolymer strategy for incorporating lipo-polymers into LNP architectures clearly has merit. Moreover, the ionizable SLO (NOx)ₙ served as a functional, and potentially superior, replacement for ALC-0315, producing higher peak transfection and greater overall expression as measured by AUC. Notably, combining the two novel POx-derived components, PMeOx-(NOx)ₙ and (NOx)ₙ, produced expression AUC values comparable to those of the industry-standard formulation, further highlighting the potential of this platform as a versatile nucleic acid delivery toolkit. Collectively, these findings establish POx-Chimera SLPs and SLOs as a promising new platform for LNP engineering and provide a foundation for the development of polymer-based alternatives to conventional PEG-lipid conjugates and ionizable lipids.

Further optimization of hydrophilic and hydrophobic block lengths, compositions, and architectures is expected to yield next-generation SLPs and SLOs capable of matching or exceeding current industry standards. The breadth of POx monomer chemistries also enables the creation of numerous ionizable SLO variants through the incorporation of different hydrophobic monomers and ionizable end groups. These next-generation materials are expected to provide unique opportunities for tuning biodistribution and transfection profiles. Ongoing work is focused on further optimization and expansion of these technologies into a broad next-generation platform for nucleic acid delivery.

## SUPPORTING INFORMATION

**Supplemental Figure 1:**
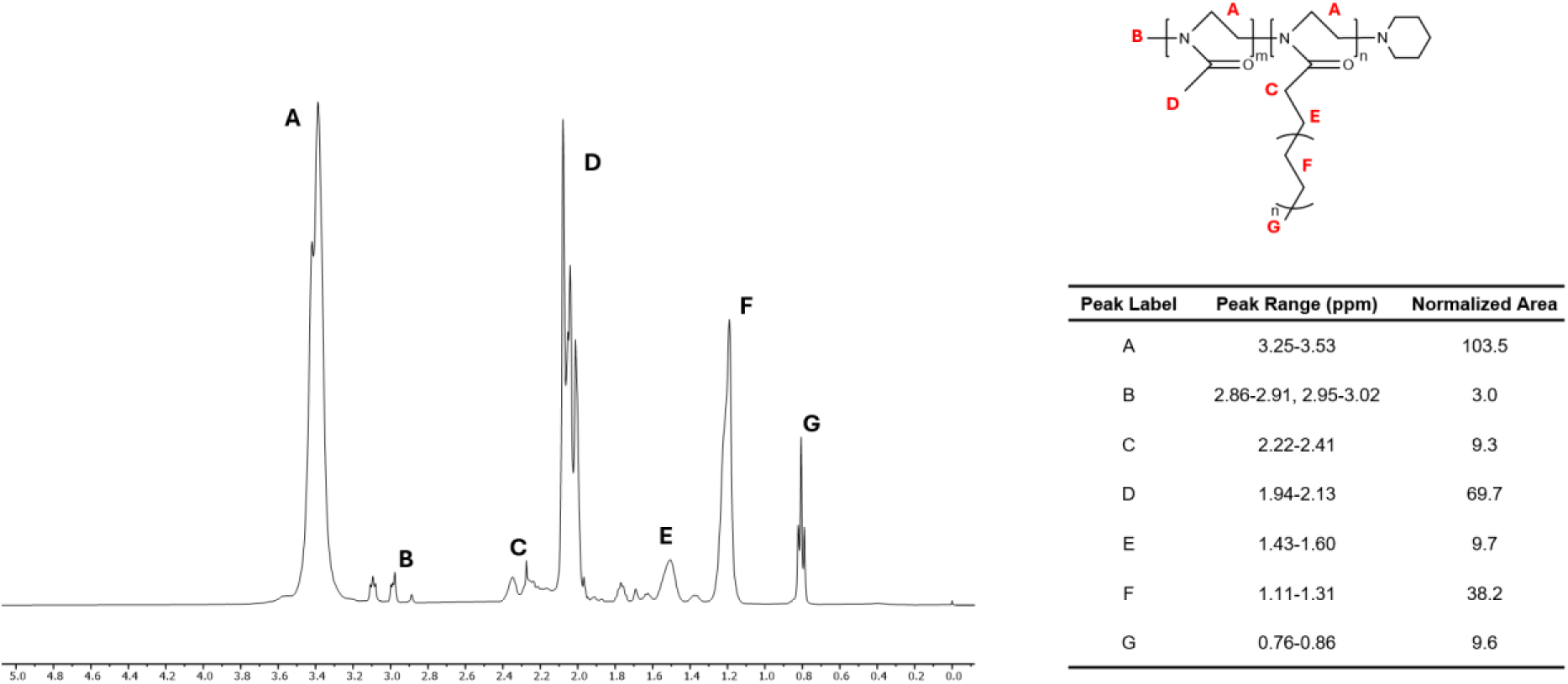
^1^HNMR spectrum and labeled molecular structure with normalized peak areas for the SLP PMeOx-(NOx)_n_

**Supplemental Figure 2:**
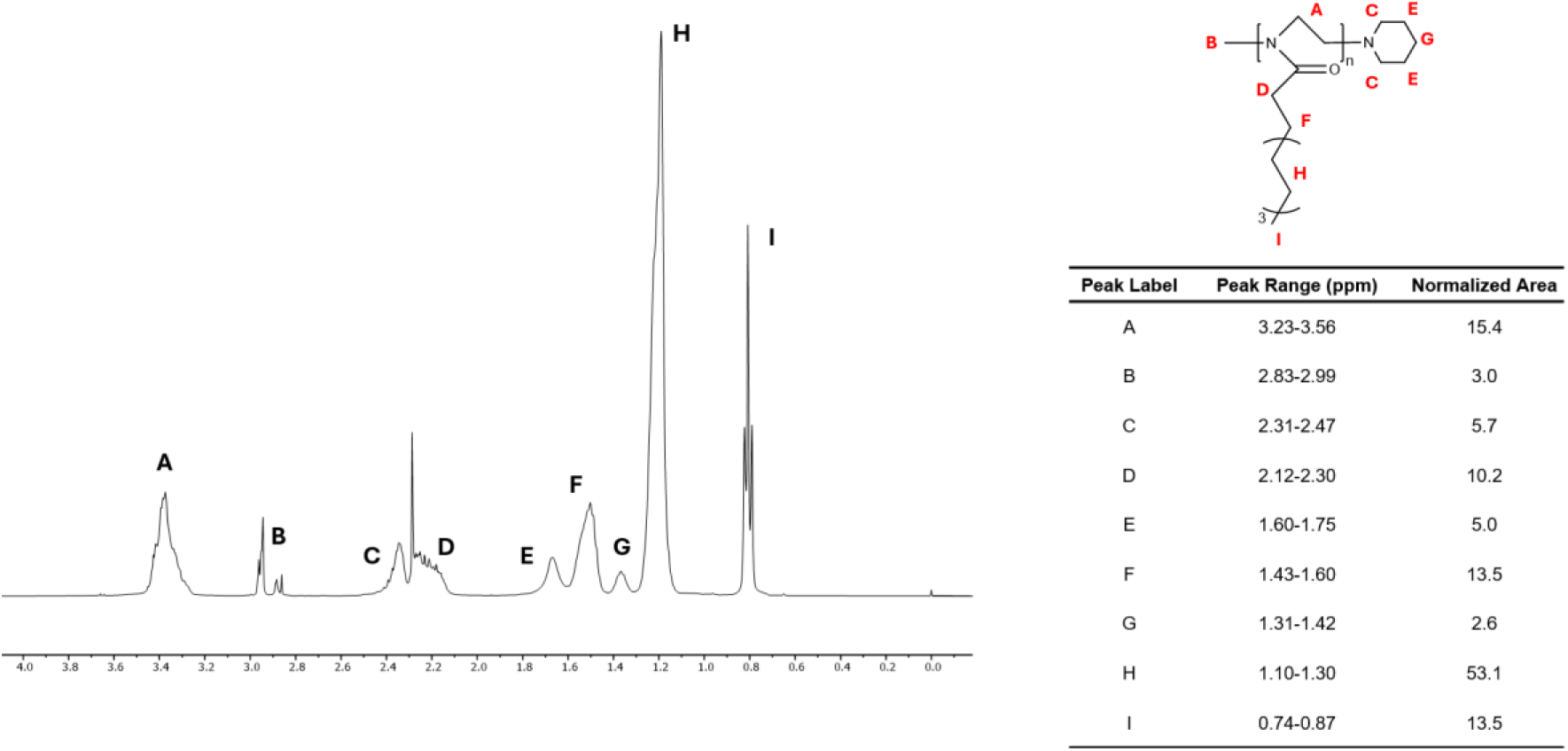
^1^HNMR spectrum and labeled molecular structure with normalized peak areas for the SLO (NOx)_n_

**Supplemental Figure 3:**
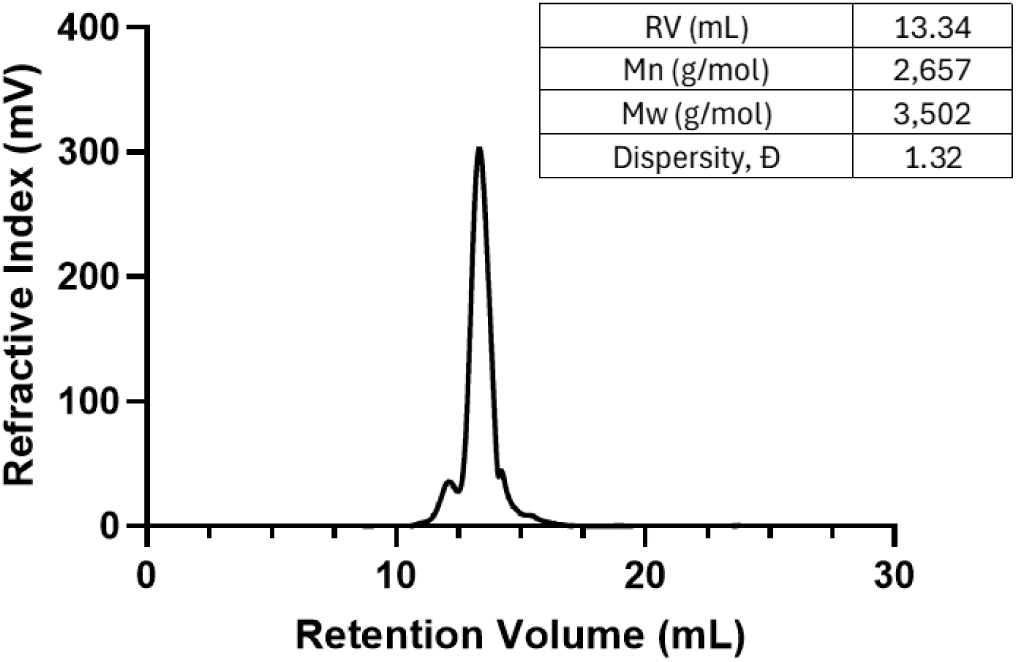
GPC elugram of PMeOx-(NOx)_n_ with M_n_, M_w_, Ð, and retention volume (RV).

**Supplemental Figure 4:**
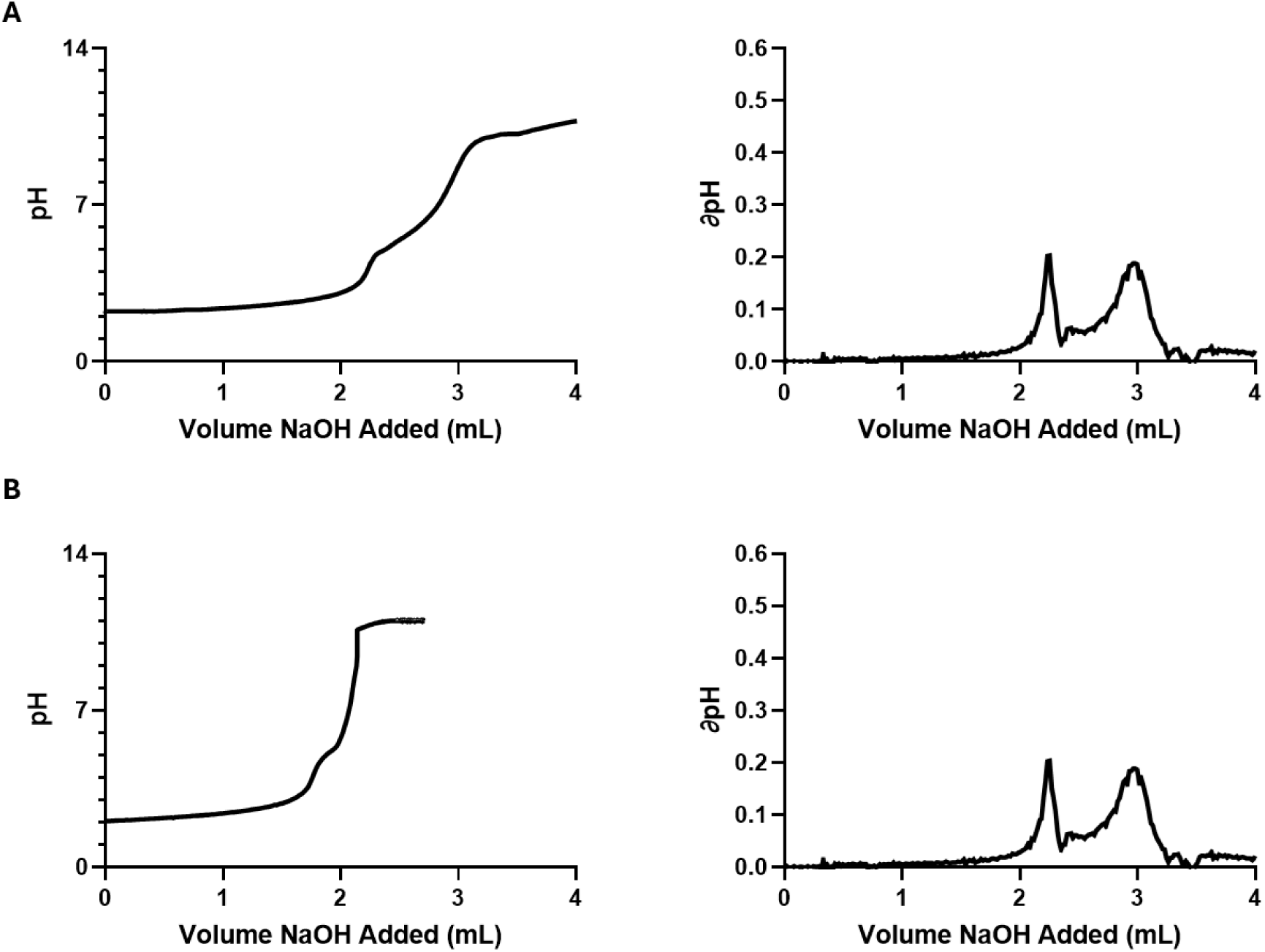
Titration and first derivative of the titration curves for **(A)** PMeOx-(NOx)_n_ and **(B)** (NOx)_n_. First equivalence point corresponds to the HCl (0.01 N) used to charge the polymer solution. Second equivalence point is polymer end-group derived ionization.

**Supplemental Table 1:**
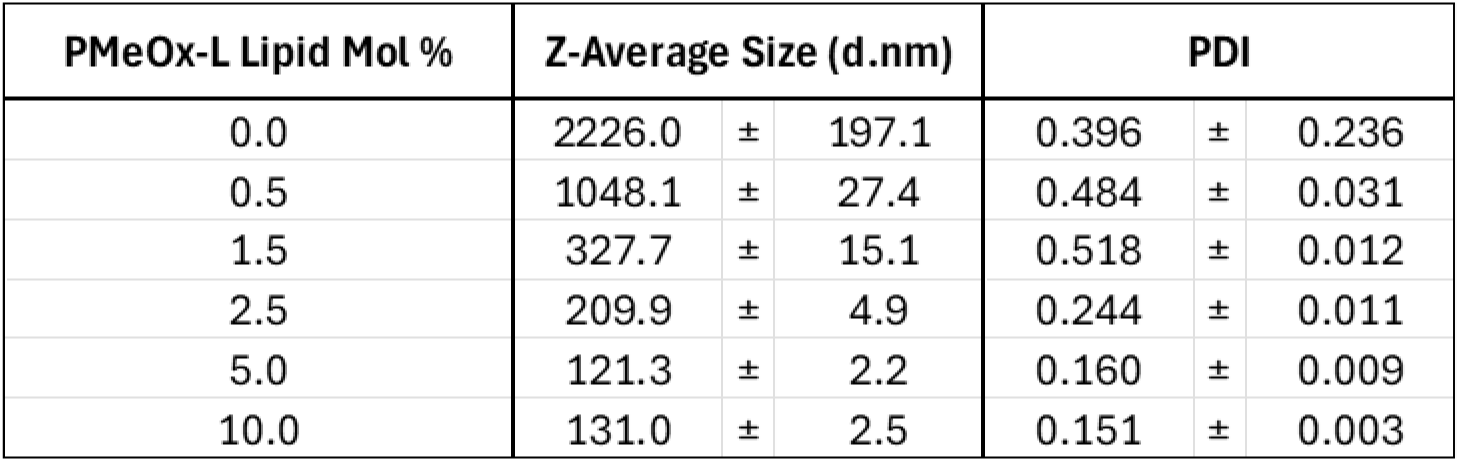
DLS size and PDI values for PMeOx-L mol % screen.

| PMeOx-L Lipid Mol % | Z-Average Size (d.nm) |  |  | PDI |  |  |
| --- | --- | --- | --- | --- | --- | --- |
| 0.0 | 2226.0 | ± | 197.1 | 0.396 | ± | 0.236 |
| 0.5 | 1048.1 | ± | 27.4 | 0.484 | ± | 0.031 |
| 1.5 | 327.7 | ± | 15.1 | 0.518 | ± | 0.012 |
| 2.5 | 209.9 | ± | 4.9 | 0.244 | ± | 0.011 |
| 5.0 | 121.3 | ± | 2.2 | 0.160 | ± | 0.009 |
| 10.0 | 131.0 | ± | 2.5 | 0.151 | ± | 0.003 |

**Supplemental Table 2:** Mol% of individual lipid components in each formulation prepared by impinged jet mixing.

| Lipid Composition |  | Mol % Components |  |  |  |
| --- | --- | --- | --- | --- | --- |
| Ionizable Lipid/Lipo-Polymer | Polymer Lipid/Lipo-Polymer | Ionizable | Polymer Lipid | Cholesterol | DSPC |
| ALC-0315 | DMG-PEG | 50.0% | 1.5% | 38.5% | 10.0% |
| ALC-0315 | PMeOx-L | 50.0% | 5.0% | 36.75% | 8.25% |
| PNOx-L | DMG-PEG | 50.0% | 1.5% | 38.5% | 10.0% |
| PNOx-L | PMeOx-L | 50.0% | 5.0% | 36.75% | 8.25% |

**Supplemental Figure 5:**
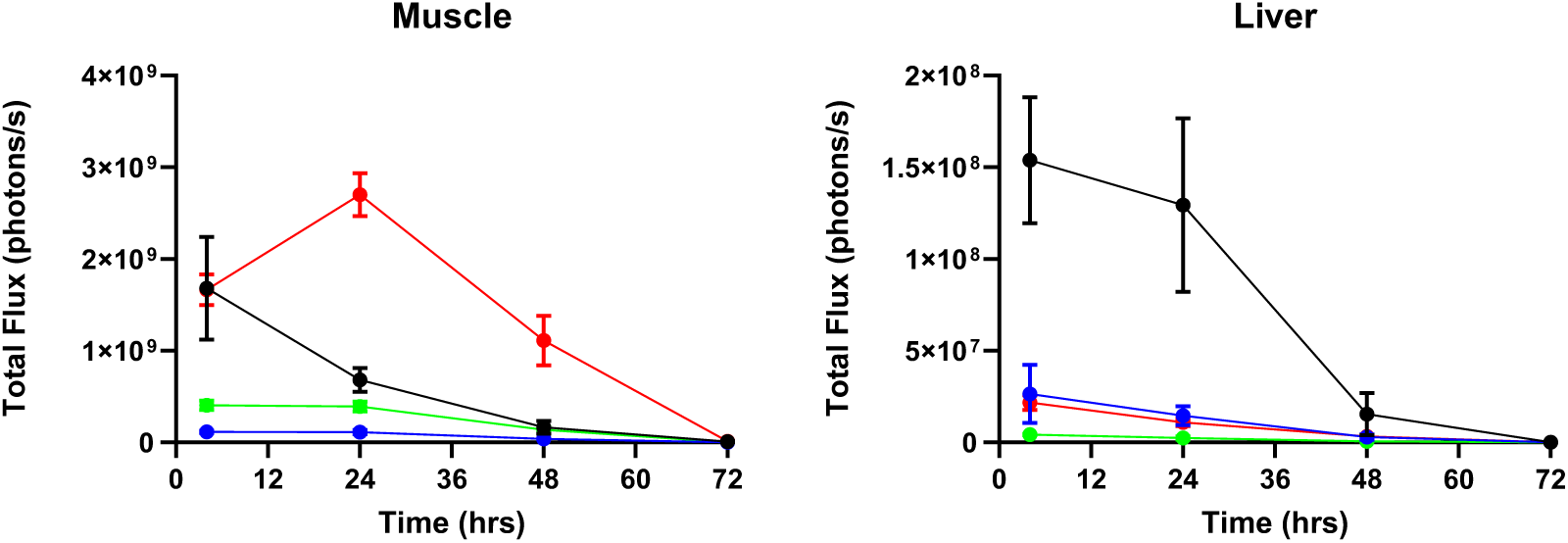
Linear y-axis presentation of IM fLuc mRNA expression from IVIS imaging.

## Materials

### Instrumentation

Mettler Toledo XS105 Dual Range Analytical Balance. Malvern OMNISEC GPC/SEC. Bruker AVANCE NEO 400 MHz Spectrometer ^1^HNMR. Malvern Zetasizer Nano ZS (Malvern Panalytical Ltd., UK). Nanodrop 2000C Spectrophotometer (ThermoFisher). Molecular Devices Spectramax M5 Plate Reader. PROMEGA 20/20 GLOMAX Luminometer. IVIS Spectrum (Revvity Inc.) in vivo Imaging System. Helix Bio Impinged Jet Mixer. Eppendorf Thermomixer R. Mettler Toledo G20S Compact Titrator.

### NOx Synthesis

Decanoic acid, ethanolamine, calcium hydride, molecular sieve, and titanium butoxide were purchased from Sigma Aldrich.

### Polymer Synthesis and Characterization

Methyl triflate was purchased from TCI Chemicals. Anhydrous acetonitrile, 2-Methyl-2-oxazoline (MeOx), and piperidine were purchased from Sigma Aldrich. HPLC grade methanol, technical grade ether, HPLC grade DMF, and Lithium Bromide (LiBr) were purchased from Fisher Scientific. House deionized DI water was used for dialysis with 1 kDa RC Spectrapor dialysis membranes used dialysis (purchased from Fisher Scientific). 0.22 µm nylon syringe filters (13 mm) were purchased from Thermofisher. Poly(methyl methacrylate) (PMMA) Gel Permeation Chromatography (GPC) standards were purchased from Sigma Aldrich. 1.0 N HCl and NaOH were purchased from Sigma Aldrich for titrations.

### Cell Culture and *in vitro* transfection

HEK293 cell line was purchased form ATCC. DMEM media (Gibco) cell culture medium and Avantor Fetal Bovine Serum (FBS) were purchased from the UNC Tissue Culture Facility. Triton X-100 reagent was purchased form Fisher Scientific. Luciferase Assay Kit was purchased from Promega. Antibiotic/Antimycotic, L-glutamine, MEM Non-essential amino acids, and 2-mercaptoethanol were purchased from Gibco.

### *In vivo* transfection

Mice (BALB/c) were purchased from Jackson Labs. D-Luciferin potassium salt for *in vivo* use from Promega (for orthotopic glioma studies). Triton X-100 reagent was purchased from Fisher Scientific. All animal studies were conducted under an approved Institutional Animal Care and Use Committee protocol at UNC Chapel Hill.

### LNP Formulation and Characterization

Cholesterol, DSPC (1,2-distearoyl-sn-glycero-3-phosphocholine), and DMG-PEG were purchased from Avanti Research. ALC-0315 was provided in collaboration with the Duke Human Vaccine Institute (DHVI). Chloroform was purchased from Fisher Scientific and 200-proof anhydrous ethanol (Supelco) was purchased from Sigma Aldrich. Sodium Acetate and Glacial Acetic Acid were purchased from Sigma Aldrich. Firefly luciferase mRNA was synthesized by and provided in collaboration with DHVI. UltraPure DNAase/RNAase free water (Invitrogen) was purchased from Fisher Scientific and 1X DPBS (Gibco) was purchased from UNC Tissue Culture Facility. Sterilized House DI water was utilized. Quant-it Ribogreen reagent was purchased from ThermoFisher Scientific. DPBS used for size, particle concentration, and zeta potential analysis was filtered with 0.2 µm nylon syringe filters (25 mm) from Fisher Scientific. Cuvettes for dynamic light scattering (DLS) (Product ZEN0040) were purchased from Malvern.

## Methods

### Monomer Synthesis

2-n-nonyl-2 oxazoline was synthesized according to the protocol described by Lübtow *et al*^23^. Briefly, a solution of decanoic acid (1.0 equivalents) was reacted with 2.0 equivalents of ethanolamine and heated to 140 °C in a Schlenk flask equipped with a dean stark trap. Reaction was allowed to proceed for ∼24 hours. Residual ethanolamine was removed under reduced pressure. 0.01 equivalent of titanium butoxide was added. The solution was stirred at 220 °C under reduced pressure. NOx was collected in the dean stark trap and dried over CaH_2_ and purified by distillation under reduced pressure. Purified product was stored over molecular sieve.

### Polymer Synthesis

POx polymers were synthesized by living cationic ring opening polymerization. Briefly, all monomers were distilled on a Schlenk line under reduced pressure/argon atmosphere to purify and remove water. Water content was confirmed below 300 ppm by Karl Fisher titration prior to use in synthesis. All glassware was dried at 120 °C overnight before use. Glassware was allowed to come to temperature under argon flow. Anhydrous acetonitrile was added to the dry schlenk flask followed by desired amount of initiator (methyl triflate). Monomer was then added to the reaction flask to achieve the desired degree of polymerization. The schlenk flask was sealed and heated at 90 °C overnight. Reaction progress was monitored by ^1^HNMR, and reaction was allowed to proceed until ∼100% monomer conversion. After full conversion and desired DP was reached, the second block (if necessary) was added. Polymers were terminated with 5-fold molar excess piperidine and allowed to react overnight at 80 °C. After termination, the reaction mixture was dissolved in small amount of methanol. Solvent was removed under reduced pressure and the polymer was dissolved again in minimal methanol. PMeOx-L was precipitated in ∼30-fold volumetric excess ice-cold diethyl ether under rapid stirring. Ether was decanted and the product dialyzed against water using 1 kDa regenerated cellulose for 3 days. PNOx-L was mixed with a small amount of toluene and then solvent removed at reduced pressure at 70 °C on a rotary evaporator. The toluene evaporation was repeated. The final PNOx-L product was a viscous liquid dissolved in chloroform and washed once with acidic water and then again with brine prior to drying to a viscous oil.

### 1HNMR Characterization

^1^HNMR spectrum were acquired using a Bruker AVANCE NEO 400 MHz Spectrometer and analyzed using the MestReNova software. Spectra were calibrated using residual solvent signal. Number-average molecular masses (M_n_) were determined via end-group analysis using ratio of initiator methyl protons to backbone and side chain protons after purification.

### GPC

GPC was performed to determine polymer molecular weight and molecular weight dispersity with a Malvern OMNISEC GPC/SEC system using a DMF (N,N-Dimethylformamide, 0.1% LiBr) mobile phase and two PolarGel-L columns (PL1117-6830) in sequence. Samples were dissolved overnight in the mobile phase at 5 mg/mL, then passed through a 13 mm 0.22 μm nylon filter into autosampler vials. Samples were analyzed with a refractive index detector during a 30 minute method with a 0.8 mL/min flow rate, 100 μL injection volume, and column and detector temperature of 25°C. A 3^rd^ order polynomial fit was used to evaluate the data using conventional calibration with PMMA standards ranging from 2,000 to 20,000 g/mol.

### Titration

Polymers were dissolved to 2 mg/mL in 0.01 N HCl and titrated on Mettler Toledo G20S Compact Titrator using 0.1 N NaOH. pKa’s were determined by plotting pH vs. volume NaOH added. Derivatives were taken and max’s found to establish inflection points and pKa’s.

### Shaker Method LNP Formulation

Screening of PMeOx-L formulation compositions was performed by the preparation of 5 μg batches of LNPs prepared based on published protocols^22^. Shaker was set to 1200 rpm and formulations were prepared at a 200 μg/mL mRNA concentration with a 3:1 aqueous to ethanol phase. *In vitro* and *in vivo* transfections were performed by diluting formulations into cell-culture media or PBS.

### Ribogreen Assay

The encapsulation efficiency and mRNA concentration was measured using a modified version of the Quant-it Ribogreen Reagent protocol. After an initial 20X dilution in DPBS, LNPs were further diluted 20X into either 1X Triton to lyse or in 1X TE Buffer supplied with the Ribogreen kit. They were each allowed to sit for 1 hour at room temperature before assaying. A 2000X dilution of the Ribogreen reagent was prepared in TE Buffer (need 100 uL per sample well. All samples were run in triplicate and standards were run in duplicate). The fLuc mRNA was diluted to 2 μg/mL and we confirmed the concentration by measuring the absorbance at 260 nm via nanodrop. A standard curve ranging from 2-100 ng RNA/mL was prepared in both TE and 0.1%. After 1 hour, each sample was further diluted 10-fold into TE buffer. 100 uL of sample or standard was to a black 96-well plate followed by 100 uL of the 2000X diluted working reagent. A blank TE and 0.1% Triton buffer control was also used as a sample for background subtraction. After 30 minutes, the fluorescence was read on a Molecular Devices SpectraMax M5 plate reader with 480 nm excitation and 520 nm emission. The encapsulation efficiency was calculated according to **Equation 1** below^10^.

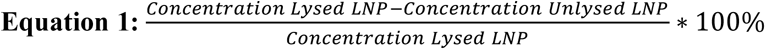

### In vitro Transfection

HEK293 cells were seeded in appropriate medium at 40,000 cells/well in a 96-well plate with n=6 wells prepared for each treatment group including blank controls. HEK293 was cultured in DMEM with 10% FBS and 1% antibiotic/antimycotic. Cells were allowed to adhere overnight. LNPs were diluted in appropriate cell culture media to 1 μg/mL total mRNA. For treatment, media was removed and 100 μL of LNPs were added to each well (100 ng mRNA/well). Cells were incubated for 24 hours. At 24 hours, media was removed and cells were lysed in 30 μL of 1X Triton and incubated with shaking at 4 °C for 1 hour. After one hour, 2 μL of each well was pipetted into a microcentrifuge tube (transparent). Sequentially and read one at a time, 50 μL of Luciferin assay reagent was added to each tube and luminescence read immediately on a ProMega GLOMAX 20/20 Luminometer^10^.

### *In vivo* Transfection

For *in vivo* studies, 1 or 3 μg of encapsulated fLuc mRNA was administered IM in 50 uL volume to N=2 or 5 female Balb/c mice (PMeOx-L screening study and optimized formulation study, respectively). Prior to injections, mice were shaved so that fur would not interfere with luminescence signals. At set times post-injection mice were injected intraperitoneally with 100 μL of 30 mg/mL D-Luciferin *in vivo* reagent. After 5 minutes, they were put under isoflurane anesthesia, and 5 minutes later luminescence was imaged on an IVIS Spectrum *in vivo* Imaging System. Images were acquired of side views and ventral views ensuring no saturated pixels were present in an image. Regions of interest (ROIs) were drawn around muscle tissue (side view) and livers (ventral view) for quantification of mRNA expression. For the PMeOx-L screening study

### LNP Manufacturing by Impinged Jet Mixing

Four-component LNPs were prepared containing ALC-0315 ionizable lipid or PNOx-L, cholesterol, DSPC, and PEG-DMG or PMeOx-L at the ratios shown in **Supplemental Table 2**. Briefly, lipids stored in chloroform were combined at necessary ratios and evaporated under nitrogen flow. Dried films were placed in a vacuum chamber until the solvent was fully evaporated. Then, the dry lipid film was solvated with ethanol to the requisite volume for mixing. Firefly luciferase mRNA was thawed at 4 °C and dissolved in a 10 mM acetate buffer at pH 4.15.

Lipids in ethanol and mRNA in buffer were combined at an amine/phosphate (N:P) ratio of 5.2:1 using a Nova Benchtop impinged jet mixer (Helix Biotech), using a flow rate ratio of 3:1 and a total flow rate of 8 mL/min. Formulated LNPs were placed in 10 kDa MWCO slide-a-lyzer devices and dialyzed against 10 mM acetate buffer, pH 4.15 for 6 hours at 4 °C; then, dialysis buffer was replaced with 1X DPBS and LNPs were dialyzed 18 hours at 4 °C. LNPs were collected and concentrated using Amicon Ultra-4 centrifugal filter units (30 kDa MWCO).

## AUTHOR INFORMATION

### Author Contributions

Jacob D. Ramsey: Conceptualization, Data Curation, Methodology, Writing, Review, and Editing, Project Administration, Funding Acquisition. Matt Haney: Data Curation, Methodology. Yuling Zhao: Data Curation, Methodology. Hallie Huttsell: Data Curation, Methodology. Alexander V. Kabanov: Conceptualization, Writing, Review, and Editing, Project Administration, Funding Acquisition.

The manuscript was written through contributions of all authors. All authors have given approval to the final version of the manuscript.

### Notes

Conflicts: JDR and AVK have a pending patent application pertaining to this matter.

## ACKNOWLEDGMENT

The authors would like to thank the Small Animal Imaging Core Facility, funded in part by NCI Cancer Center Support Grant P30CA016086, and the NIH S10 Share Instrumentation Grant (NMR) S10OD026951 for supporting this work. We would like to acknowledge Fadi Marayati and the Duke Human Vaccine Institute for their collaboration in mRNA synthesis and lipid selection.

## ABBREVIATIONS

LNP: Lipid Nanoparticle
PEG: poly(ethylene-glycol)
PEG: Poly(2-oxazoline)
SLP: Synthetic lipid polymer
SLO: Synthetic lipid oligomer
PMeOx: Poly(2-methyl-2-oxazoline)
PNOx: Poly(2-nonyl-2-oxazoline)
GPC: Gel permeation chromatography
fLuc: Firefly luciferase
DLS: Dynamic light scattering
IM: Intramuscular
AUC: Area under the curve

## REFERENCES

(1) Hou, X.; Zaks, T.; Langer, R.; Dong, Y. Lipid Nanoparticles for mRNA Delivery. Nat Rev Mater 2021, 6 (12), 1078–1094. 10.1038/s41578-021-00358-0

(2) Jeong, M.; Lee, Y.; Park, J.; Jung, H.; Lee, H. Lipid Nanoparticles (LNPs) for *in Vivo* RNA Delivery and Their Breakthrough Technology for Future Applications. Advanced Drug Delivery Reviews 2023, 200, 114990. 10.1016/j.addr.2023.114990

(3) Kon, E.; Ad-El, N.; Hazan-Halevy, I.; Stotsky-Oterin, L.; Peer, D. Targeting Cancer with mRNA–Lipid Nanoparticles: Key Considerations and Future Prospects. Nat Rev Clin Oncol 2023, 20 (11), 739–754. 10.1038/s41571-023-00811-9

(4) Samaridou, E.; Heyes, J.; Lutwyche, P. Lipid Nanoparticles for Nucleic Acid Delivery: Current Perspectives. Advanced Drug Delivery Reviews 2020, 154*–*155, 37–63. 10.1016/j.addr.2020.06.002

(5) Cheng, Q.; Wei, T.; Farbiak, L.; Johnson, L. T.; Dilliard, S. A.; Siegwart, D. J. Selective Organ Targeting (SORT) Nanoparticles for Tissue-Specific mRNA Delivery and CRISPR– Cas Gene Editing. Nat. Nanotechnol. 2020, 15 (4), 313–320. 10.1038/s41565-020-0669-6

(6) Ren, Y.; Lin, L.; Abdallah, M.; Zhu, X.; Liu, H.; Fabb, S. A.; Payne, T. J.; Pouton, C. W.; Johnston, A. P. R.; Trevaskis, N. L. Impact of Ionizable Lipid Type on the Pharmacokinetics and Biodistribution of mRNA-Lipid Nanoparticles after Intravenous and Subcutaneous Injection. J Control Release 2025, 384, 113945. 10.1016/j.jconrel.2025.113945

(7) Kim, M.; Song, E. S.; Chen, J. C.; Chatterjee, S.; Sun, Y.; Lee, S. M.; Wu, S.; Patel, P.; Tian, Z.; Kantor, A.; Wustman, B. A.; Lockhart, D. J.; Siegwart, D. J. Dual SORT LNPs for Multi-Organ Base Editing. Nat Biotechnol 2026, 44 (4), 578–586. 10.1038/s41587-025-02675-z

(8) Swingle, K. L.; Hamilton, A. G.; Safford, H. C.; Geisler, H. C.; Thatte, A. S.; Palanki, R.; Murray, A. M.; Han, E. L.; Mukalel, A. J.; Han, X.; Joseph, R. A.; Ghalsasi, A. A.; Alameh, M.-G.; Weissman, D.; Mitchell, M. J. Placenta-Tropic VEGF mRNA Lipid Nanoparticles Ameliorate Murine Pre-Eclampsia. Nature 2025, 637 (8045), 412–421. 10.1038/s41586-024-08291-2

(9) Vasudevan, A.; Jozić, A.; Curtis, A. G.; Bodi, E.; Ryals, R. C.; Sahay, G. Lipid Nanoparticle-Mediated Intracameral mRNA Delivery Facilitates Gene Expression and Editing in the Anterior Chamber of the Eye. J Control Release 2025, 379, 1022–1028. 10.1016/j.jconrel.2025.01.068

(10) Basham, C.; Haney, M.; Zhao, Y.; Lukyanov, K. A.; Kim, K.; Baysal, A.; Hutsell, H.; Kabanov, A. V.; Ramsey, J. D. PEG-Free Tunable Poly(2-Oxazoline) Lipids Modulate LNP Biodistribution and Expression In Vivo after Intramuscular Administration. bioRxiv 2025, 2025.06.05.657891. 10.1101/2025.06.05.657891

(11) Holick, C. T.; Klein, T.; Mehnert, C.; Adermann, F.; Anufriev, I.; Streiber, M.; Harder, L.; Traeger, A.; Hoeppener, S.; Franke, C.; Nischang, I.; Schubert, S.; Schubert, U. S. Poly(2-ethyl-2-oxazoline) (POx) as Poly(Ethylene Glycol) (PEG)-Lipid Substitute for Lipid Nanoparticle Formulations. Small 2025, 21 (16), 2411354. 10.1002/smll.202411354

(12) Golba, B.; Zhong, Z.; Romio, M.; Almey, R.; Deforce, D.; Dhaenens, M.; Sanders, N. N.; Benetti, E. M.; De Geest, B. G. Cyclic Poly(2-Methyl-2-Oxazoline)-Lipid Conjugates Are Good Alternatives to Poly(Ethylene Glycol)-Lipids for Lipid Nanoparticle mRNA Formulation. Biomacromolecules 2025, 26 (3), 1816–1825. 10.1021/acs.biomac.4c01587

(13) He, X.; Payne, T. J.; Takanashi, A.; Fang, Y.; Kerai, S. D.; Morrow, J. P.; Al-Wassiti, H.; Pouton, C. W.; Kempe, K. Tailored Monoacyl Poly(2-Oxazoline)- and Poly(2-Oxazine)-Lipids as PEG-Lipid Alternatives for Stabilization and Delivery of mRNA-Lipid Nanoparticles. Biomacromolecules 2024, 25 (7), 4591–4603. 10.1021/acs.biomac.4c00651

(14) Moreadith, R. W.; Kim, W.; Smith, K.; Yoon, K.; Weimer, R.; Fang, Z. Overcoming the PEG Dilemma with Poly(2-Ethyl-2-Oxazoline) Lipids in Lipid Nanoparticle Formulations. European Polymer Journal 2025, 241, 114392. 10.1016/j.eurpolymj.2025.114392

(15) Sanchez, A. J. D. S.; Loughrey, D.; Echeverri, E. S.; Huayamares, S. G.; Radmand, A.; Paunovska, K.; Hatit, M.; Tiegreen, K. E.; Santangelo, P. J.; Dahlman, J. E. Substituting Poly(Ethylene Glycol) Lipids with Poly(2-Ethyl-2-Oxazoline) Lipids Improves Lipid Nanoparticle Repeat Dosing. Adv Healthc Mater 2024, 13 (17), e2304033. 10.1002/adhm.202304033

(16) van Zyl, D. G.; Mendes, L. P.; Semper, R. P.; Rueckert, C.; Baumhof, P. Poly(2-Methyl-2-Oxazoline) as a Polyethylene Glycol Alternative for Lipid Nanoparticle Formulation. *Front*. Drug Deliv. 2024, 4. 10.3389/fddev.2024.1383038

(17) Guerrini, G.; Gioria, S.; Sauer, A. V.; Lucchesi, S.; Montagnani, F.; Pastore, G.; Ciabattini, A.; Medaglini, D.; Calzolai, L. Monitoring Anti-PEG Antibodies Level upon Repeated Lipid Nanoparticle-Based COVID-19 Vaccine Administration. Int J Mol Sci 2022, 23 (16), 8838. 10.3390/ijms23168838

(18) Zhang, P.; Sun, F.; Liu, S.; Jiang, S. Anti-PEG Antibodies in the Clinic: Current Issues and beyond PEGylation. Journal of Controlled Release 2016, 244, 184–193. 10.1016/j.jconrel.2016.06.040

(19) McSweeney, M. D.; Price, L. S. L.; Wessler, T.; Ciociola, E. C.; Herity, L. B.; Piscitelli, J. A.; DeWalle, A. C.; Harris, T. N.; Chan, A. K. P.; Saw, R. S.; Hu, P.; Jennette, J. C.; Forest, M. G.; Cao, Y.; Montgomery, S. A.; Zamboni, W. C.; Lai, S. K. Overcoming Anti-PEG Antibody Mediated Accelerated Blood Clearance of PEGylated Liposomes by Pre-Infusion with High Molecular Weight Free PEG. Journal of Controlled Release 2019, 311*–*312, 138–146. 10.1016/j.jconrel.2019.08.017

(20) Ju, Y.; Kent, S. J. Balancing Stealth and Targeting to Improve Nanomedicine Efficacy. Nat Nanotechnol 2025, 20 (5), 576–579. 10.1038/s41565-025-01926-z

(21) Dobrovolskaia, M. A. Better the Devil You Know than the Devil You Don’t — PEG Challenges in Nanomedicine. Nat. Nanotechnol. 2025, 20 (5), 580–583. 10.1038/s41565-025-01925-0

(22) Yoo, S.; Han, E. L.; Murray, A. M.; Kim, D.; Bang, E.-K.; Mitchell, M. J. High-Throughput Formulation Using Fast mRNA Lipid Nanoparticle Assembly via Simple Thermomixing (HTF-FLASH). Nano Lett 2025, 25 (40), 14695–14703. 10.1021/acs.nanolett.5c03793

(23) Lübtow, M. M.; Keßler, L.; Appelt-Menzel, A.; Lorson, T.; Gangloff, N.; Kirsch, M.; Dahms, S.; Luxenhofer, R. More Is Sometimes Less: Curcumin and Paclitaxel Formulations Using Poly(2-Oxazoline) and Poly(2-Oxazine)-Based Amphiphiles Bearing Linear and Branched C9 Side Chains. Macromol Biosci 2018, 18 (11), e1800155. 10.1002/mabi.201800155

